# Quantitative proteomics reveals differential extracellular vesicle cargo from M1 and M2 monocyte-derived human macrophages

**DOI:** 10.1101/2024.09.17.613483

**Authors:** Paschalia Pantazi, Toby Clements, Harriet T Parsons, Myrsini Kaforou, Kate J Heesom, Phillip R Bennett, Seth Guller, Vikki M Abrahams, Beth Holder

## Abstract

Extracellular vesicles (EVs) mediate intercellular communication by carrying molecular cargo that facilitate diverse physiological processes. Macrophages, playing central roles in immune responses, release EVs that modulate various cellular functions. Given the distinct roles of M1 and M2 macrophage states, understanding the proteomic profiles of their EVs is important for elucidation of EV-mediated signalling and identifying potential biomarkers for diseases involving macrophage polarisation. We employed quantitative proteomics combined with bioinformatics to characterise the proteomic profile of EVs released by M1 and M2 monocyte-derived macrophages. We identified 1,731 proteins in M1/M2 EVs, 132 of which were significantly differentially between M1 and M2. Proteomic data, together with pathway analysis, found that M1/M2 macrophage EV cargo relate to cellular source, and may play roles in shaping immune responses, with M1 EV cargo associated with promotion of pro-inflammatory and antiviral functions, while M2 EV cargo associated with immune regulation and tissue repair. M1 EV cargo was associated with cytokine/chemokine signalling pathways, DNA damage, methylation, and oxidative stress. M2 EV cargo were associated with macrophage alternative-activation signalling pathways, antigen presentation, and lipid metabolism. We also report that macrophage EVs carry metallothioneins, and other related proteins involved in response to metals and oxidative stress.

## INTRODUCTION

Extracellular vesicles (EVs) have emerged as key mediators of intercellular communication, playing important roles in physiological and pathological processes across diverse biological systems. These nanoscale membrane-encapsulated structures, released by cells into the extracellular environment, carry a range of cargo, including proteins, nucleic acids, and lipids, thereby facilitating the exchange of molecular information between cells. Macrophages, as efficient antigen presenting cells found throughout adult tissues, exhibit a diverse array of functions, including pathogen defence, wound healing, and immune cell modulation. Many of these functions may involve EVs as key mediators, with macrophage EVs shown to regulate fibroblast proliferation [1], mediate endothelial cell activation [2], modulate insulin sensitivity [3], and traffic pathogen elements to uninfected cells [4,5], among many other reported actions [6].

Proteomic profiling of EV cargo can provide useful data for understanding both autocrine and paracrine EV messaging. Several proteomic studies exist relating to immune cell EVs, but have often focused on cell lines or cells derived from murine bone marrow (such as bone marrow-derived macrophages) with fewer studies on primary human immune cell EV proteomic cargo, due to limitations with isolating sufficient numbers of cells for EV generation. Previous proteomic analysis of EVs from primary human monocyte-derived macrophages have identified modification of cargo following in vitro Influenza A virus infection [7] and following β-glucan stimulation [8].

Macrophages exist in several phenotypic states essential for orchestrating different immune responses, and for support of homeostasis. Among these, M1 and M2 macrophages represent contrasting functional profiles; M1 macrophages exhibit pro-inflammatory and microbicidal activities, while M2 macrophages display anti-inflammatory and tissue repair functions [9]. The M1/M2 paradigm has some limitations, but is the most widely utilised model of macrophage function. Thus, data on the EVs from these cells are highly useful for the study of EV messaging between macrophages and recipient cells in health and disease. We have previously reported the non-coding RNA cargo of M1/M2 macrophage EVs [10]. Here we have performed quantitative proteomics on EVs released from these two macrophage polarisation states. This dataset provides a foundation for future research into macrophage EV biology, and offers potential biomarkers for future studies of the many diseases associated with inappropriate M1 macrophage polarisation.

## MATERIALS AND METHODS

### Generation of M1 and M2 monocyte-derived macrophages

Blood was acquired from Research Donors, a HTA licenced and ISO 9001 2015 certified company with Research Ethics Committee (REC) approval (Reference: 20/LO/0325). M1/M2 macrophages were generated as before[10]. In brief, peripheral blood from non-fasted healthy female donors (21-40 years) was collected in sodium-heparin tubes and transported to the lab within 6 hours. Platelets were depleted by centrifugation, followed by PBMC isolation using Histopaque-1077, with additional steps to deplete platelets. PBMC were seeded in serum-free X-VIVO 10 media with 1% penicillin-streptomycin. Monocytes adhered for 1 hour, and non-adherent/weakly attached cells were removed through vigorous washing. M1 cells were induced with 20ng/mL of granulocyte-macrophage colony-stimulating factor (GM-CSF, Cat. No. 572904, BioLegend), while M2 macrophages were induced with 20 ng/mL of macrophage–colony-stimulating factor (M-CSF, Cat. No. 574804, BioLegend). After 6 days, 50% additional fresh media with M-CSF/GM-CSF was added. On day 7, to achieve full polarization, M1 cultures were exposed to 20 ng/mL of IFN-γ (Cat. No. 300-02, Peprotech) and 20 ng/mL LPS (Cat. No. L2137, Sigma). M2 cells were treated with 20 ng/mL IL-4 (Cat. No. 200-04, Peprotech) and 20 ng/mL IL-13 (Cat. No. 200-13, Peprotech). These final treatments were 48 hours, with supplemented media refreshed after 24 hours.

### M1/M2 macrophage extracellular vesicle isolation

EVs were isolated from M1/M2 macrophages as before [10]. In brief, following generation of fully polarised M1/M2 macrophages, fresh serum-free X-VIVO 10 media with 1% penicillin-streptomycin was added for a 24-hour EV generation period. Conditioned media was centrifuged at 300 xG for 5 mins, at 1000 xG for 10 mins, and then concentrated to 500 µL using Vivaspin 15R Hydrosart 30,000 MWCO columns (Cat. No. FIL8452, Sartorius). This 500 µL was loaded onto 70nm qEV Original size exclusion chromatography columns (Cat. No. SP1, IZON) in an Automatic Fraction Collector (IZON) for EV isolation into DPBS. Following a 2.8ml void volume, fractions of 500 µL were collected.

### Nanoparticle tracking analysis (NTA)

EVs were counted and sized by Nanoparticle Tracking Analysis (NTA) using the ZetaView PMX 120 S (Particle Metrix). The instrument utilized the standard NTA cell assembly and was controlled by the ZetaView 8.05.12 SP2 software. Prior to sample readings, the instrument was calibrated using polymer beads with a consistent size of 100nm (Cat. No. 3100A, Thermo Fisher). Samples were appropriately diluted in 1 mL PBS, ensuring a concentration within the recommended range of 50-200 particles/frame (dilutions ranging from 1:100 to 1:250). Each sample was subjected to a 21-second video capture at 11 different positions, with two readings at each position. Following automated analysis and removal of outliers, the software calculated the size and concentration of the samples. Pre-acquisition parameters were established at a temperature of 23°C, a sensitivity of 75, a frame rate of 30 frames per second, a shutter speed of 100, and a laser pulse duration equivalent to the shutter duration. Post-acquisition parameters included a minimum brightness of 30, a maximum size of 1000 pixels, a minimum size of 10 pixels, and a trace length of 15.

### Electron microscopy

Following establishment of EV elution patterns by SEC, EV-enriched fractions (1-3) were combined and concentrated using Amicon Ultra 0.5mL 30kDa cut-off spin columns (Cat. No. 10012584, Fisher Scientific). 300 mesh continuous carbon support copper grids (Cat. No. AGG2300C, Agar Scientific) were glow discharged using a Fischione NanoClean Model 1070 instrument. Six microliters of each concentrated sample were applied directly on the grids for 5 min followed by 2% uranyl acetate for 45 sec. The grids were then air dried and imaged using a Tecnai T12 transmission electron microscope at 120 kVolt.

### Protein isolation and quantification

Protein was isolated from the concentrated EVs by addition of RIPA buffer to 1X concentration (Cat. No. 10010263, Cayman Chemical) supplemented with protease (Cat. No. 04693124001, Roche) and phosphatase (Cat. No. 4906837001, Sigma) inhibitors. Lysates were incubated on ice for 1 h and then the three EV containing fractions of each sample were pooled together and concentrated using the Vivacon® 500, 2 kDa cut-off Hydrosart columns (Cat. No. VN01H91, Sartorius). Protein was quantified by microBCA assay (Cat. No. 23227, Thermo Fisher), read at 562 nm on a Nanodrop 2000 spectrophotometer (Thermo Scientific).

### TMT Labelling and High pH reversed phase chromatography

Aliquots of 15 µg of each sample were digested with trypsin (1.25 µg trypsin, 37°C, overnight), labelled with Tandem Mass Tag (TMT) 10-plex reagents according to the manufacturer’s protocol (Thermo Fisher Scientific) and the labelled samples pooled. The pooled sample was desalted using a SepPak cartridge according to the manufacturer’s instructions (Waters). Eluate from the SepPak cartridge was evaporated to dryness and resuspended in buffer A (20 mM ammonium hydroxide, pH 10) prior to fractionation by high pH reversed-phase chromatography using an Ultimate 3000 liquid chromatography system (Thermo Fisher Scientific). In brief, the sample was loaded onto an XBridge BEH C18 Column (130Å, 3.5 µm, 2.1 mm X 150 mm, Waters, UK) in buffer A and peptides eluted with agcan increasing gradient of buffer B (20 mM Ammonium Hydroxide in acetonitrile, pH 10) from 0-95% over 60 minutes. The resulting fractions (5 in total) were evaporated to dryness and resuspended in 1% formic acid prior to analysis by nano-LC MSMS using an Orbitrap Fusion Lumos mass spectrometer (Thermo Scientific).

### Nano-LC Mass Spectrometry

High pH RP fractions were further fractionated using an Ultimate 3000 nano-LC system in line with an Orbitrap Fusion Lumos mass spectrometer (Thermo Scientific). In brief, peptides in 1% (vol/vol) formic acid were injected onto an Acclaim PepMap C18 nano-trap column (Thermo Scientific). After washing with 0.5% (vol/vol) acetonitrile 0.1% (vol/vol) formic acid peptides were resolved on a 250 mm × 75 μm Acclaim PepMap C18 reverse phase analytical column (Thermo Scientific) over a 150 min organic gradient, using 7 gradient segments (1-6% solvent B over 1 min., 6-15% B over 58 min., 15-32% B over 58 min., 32-40% B over 5 min., 40-90% B over 1 min., held at 90% B for 6 min and then reduced to 1% B over 1 min.) with a flow rate of 300 nl/min. Solvent A was 0.1% formic acid and Solvent B was aqueous 80% acetonitrile in 0.1% formic acid. Peptides were ionized by nano-electrospray ionization at 2.0 kV using a stainless-steel emitter with an internal diameter of 30μm (Thermo Scientific) and a capillary temperature of 300°C.

All spectra were acquired using an Orbitrap Fusion Lumos mass spectrometer controlled by Xcalibur 3.0 software (Thermo Scientific) and operated in data-dependent acquisition mode using an SPS-MS3 workflow. FTMS1 spectra were collected at a resolution of 120,000, with an automatic gain control (AGC) target of 200,000 and a max injection time of 50 ms. Precursors were filtered with an intensity threshold of 5,000, according to charge state (to include charge states 2-7) and with monoisotopic peak determination set to Peptide. Previously interrogated precursors were excluded using a dynamic window (60s +/-10ppm). The MS2 precursors were isolated with a quadrupole isolation window of 0.7m/z. ITMS2 spectra were collected with an AGC target of 10,000, max injection time of 70ms and CID collision energy of 35%.

For FTMS3 analysis, the Orbitrap was operated at 50,000 resolution with an AGC target of 50,000 and a max injection time of 105 ms. Precursors were fragmented by high energy collision dissociation at a normalised collision energy of 60% to ensure maximal TMT reporter ion yield. Synchronous Precursor Selection (SPS) was enabled to include up to 10 MS2 fragment ions in the FTMS3 scan.

### Proteomic Analysis

The raw data files were processed and quantified using Proteome Discoverer software v3.1 (Thermo Scientific) and searched against the UniProt Human database (reviewed, canonical proteins and reviewed isoforms, downloaded January 2024: 42,508 entries) using the SEQUEST HT algorithm. Peptide precursor mass tolerance was set at 10ppm, and MS/MS tolerance was set at 0.6 Da. Search criteria included oxidation of methionine (+15.995Da), acetylation of the protein N-terminus (+42.011Da) and Methionine loss plus acetylation of the protein N-terminus (−89.03Da) as variable modifications and carbamidomethylation of cysteine (+57.021Da) and the addition of the TMT mass tag (+229.163 Da) to peptide N-termini and lysine as fixed modifications. Searches were performed with full tryptic digestion and a maximum of 2 missed cleavages were allowed. The reverse database search option was enabled and all data was filtered to satisfy false discovery rate (FDR) of 5%.

### Comparison of M1 and M2 EV protein cargo

Differential abundance analysis was performed using Lava 2.0 (https://github.com/tempeparsons/Lava). Briefly, this measures the log2 difference between two groups and performs and independent T-test using the numpy (https://numpy.org/) and scipy packages (https://docs.scipy.org/doc/scipy/reference/generated/scipy.stats.ttest_ind_from_stats.html, parameters set as equal_var=True, alternative=’two-sided’) in the Python 3.10 programming language. Protein abundance was measured using scaled abundance values from PD 3.1. For volcano plots, a linear fold-change threshold was set at 2 or Log2(1). Log2(fold change) was plotted against negative transformed Log10(p-value), with the y-axis re-labelled to show the actual p-value. Data points were sized according to abundance and protein identities based on a single peptide (’low pep count’) were coloured grey

### Bioinformatic analysis

Prior to further analysis, additional isoforms were removed from the dataset, to aid interpretation and downstream pathway analysis. The lists of significantly upregulated proteins in M1 or M2 EVs (peptide count>1, FC>2, adjusted p value<0.05) were subjected to separate pathway analysis using Ingenuity Pathway Analysis (IPA; QIAGEN Inc., https://www.qiagenbioinformatics.com/products/ingenuity-pathway-analysis), which predicts directionality of pathways through knowledge of molecular functions. The resulting z scores indicate degree of change between the groups, with positive and negative z-scores indicating activation and repression of a pathway, respectively.

## RESULTS AND DISCUSSION

### Isolation of EVs from M1/M2 mo-macrophages

M1 and M2 macrophages were generated from monocytes of healthy donors, in the presence of GM-CSF followed by IFN-γ and LPS, or M-CSF followed by IL-13 and IL-4, respectively (Figure 1A), and as previously described [10]. EVs were successfully isolated from M1 and M2 primary macrophages using size exclusion chromatography, and were enriched in fractions 1-3 following void, as measured by nanoparticle tracking analysis (NTA; Fig 1B), which were pooled for subsequent analysis. EV concentration per million starting PBMC was 9.8×10^7^ for M1 and 5.0×10^7^ for M2. EVs exhibited the expected size profile of EVs (Fig 1C). The median size of the EVs was 161.4nm for M1 and 155.5 for M2 (Fig 1E), and the mean size was 174.2nm for M1 and 171.6nm for M2 EVs (Fig 1F). We have previously reported additional characterisation of our M1/M2 macrophage EVs, including morphology by transmission electron microscopy, and EV marker detection by western blotting and ELISA [10].

**Figure 1.**
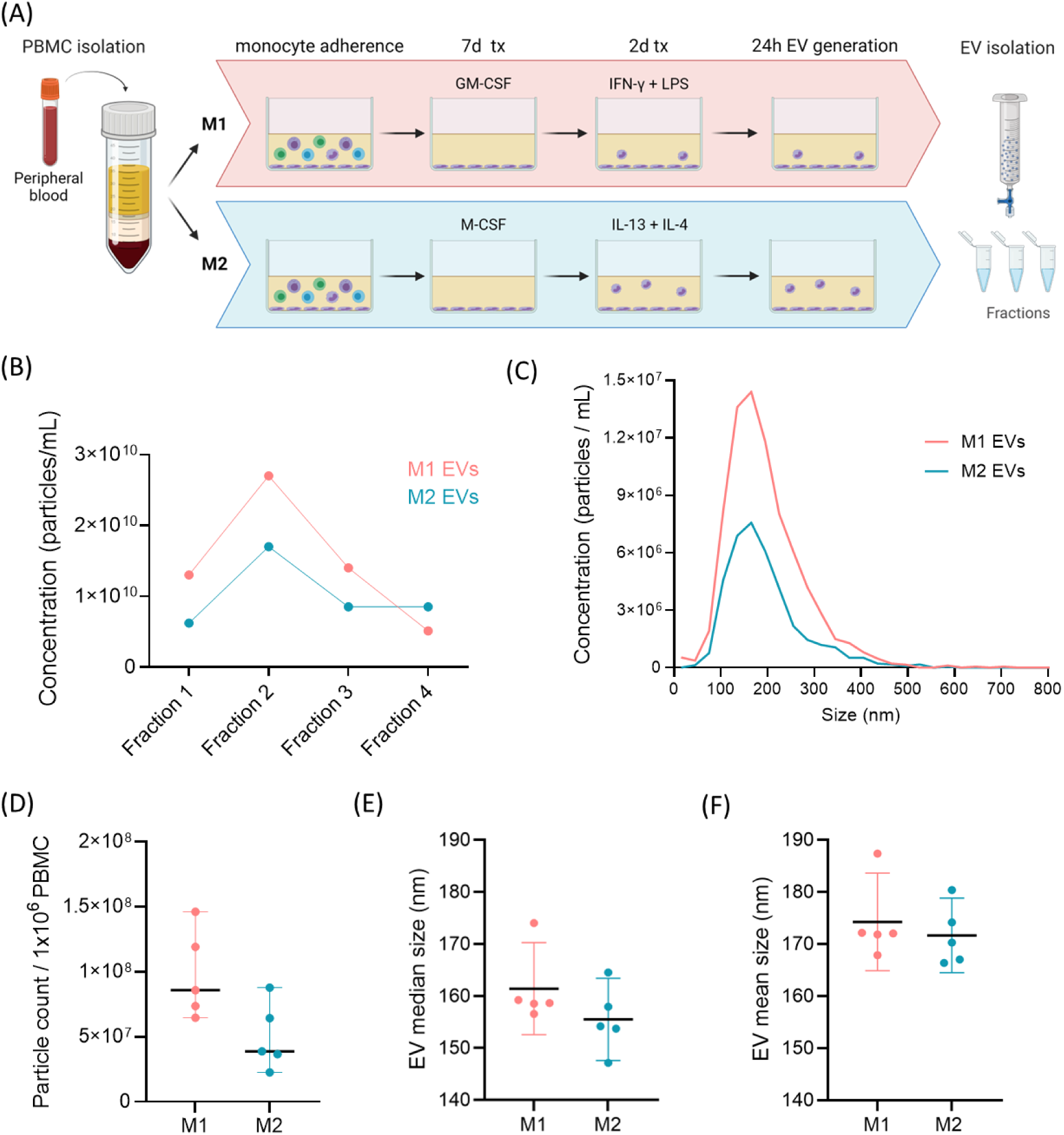
Isolation and characterisation of M1/M2 macrophage extracellular vesicles. A) Schematic representation of EV generation from M1 and M2 monocyte-derived macrophages. B) Representative particle concentration of EV-containing fractions isolated by size exclusion chromatography, following void, measured by NTA. C) Representative size distribution of M1 and M2 EVs (pooled fractions 1-3) D) Total particle count measured by NTA in fractions 1-3, per million PBMC seeded, mean +95% CI, n=5 E) Median size of EVs from M1 and M2 cells, measured by NTA, mean +95% CI, n=5.

### Protein cargo of extracellular vesicles from M1 and M2 macrophages

Using TMT-based quantitative mass spectrometry, we identified 2,129 proteins in EVs from monocyte-derived macrophages (Supplementary table 1, tab 1). 307 proteins were present as more than one isoform, representing a total of 705 identified proteins. Removal of the additional isoforms left us with a final list of 1,731 proteins, which were taken forward for further analysis (Supplemental table 1, tab 2). This list contained 99 of the 100 most commonly reported proteins on the Vesiclepedia repository (accessed 18^th^ March 2024) [11], reinforcing the reliability of our workflow. A visual summary of proteins found in M1/M2 macrophage EVs that are commonly reported as EV cargo/markers, is provided in Figure 2A. This included a range of EV surface proteins, such as heat shock proteins (HSPs), tetraspanins (CD9, CD81, CD9, etc), several annexins, integrins, selectins, and Human Leukocyte Antigens (HLAs). EV markers expected to be located in the lumen, included members of the ESCRT (endosomal sorting complexes required for transport) pathway (e.g. TSG101 and VSPs), cytoskeletal proteins, enzymes, and proteins involved in membrane transport, such as Rab proteins, flotillins and the transferrin receptor. In terms of proteins relevant to macrophage function, monocyte-derived macrophage EVs carried both MHC class II and associated co-stimulatory molecules, including CD80 and CD86 (Figure 2B). Of the pattern recognition receptors (PRRs), which are involved in responses to pathogen-associated patterns (PAMPs), a restricted number of toll-like receptors (TLRs) were seen, with only TLR2 and TLR9 being detectable. Many C-type lectin receptors were present, including common macrophage markers CD206 (MRC1) and CD209 (DC-SIGN), and several scavenger receptors, including CD14, CD36 and MSR1. We did not detect other PRR classes, i.e. Nucleotide-binding Oligomerization Domain (NOD)-like Receptors (NLRs), and Retinoic Acid-inducible Gene I (RIG-I)-like Receptors (RLRs).

**Figure 2.**
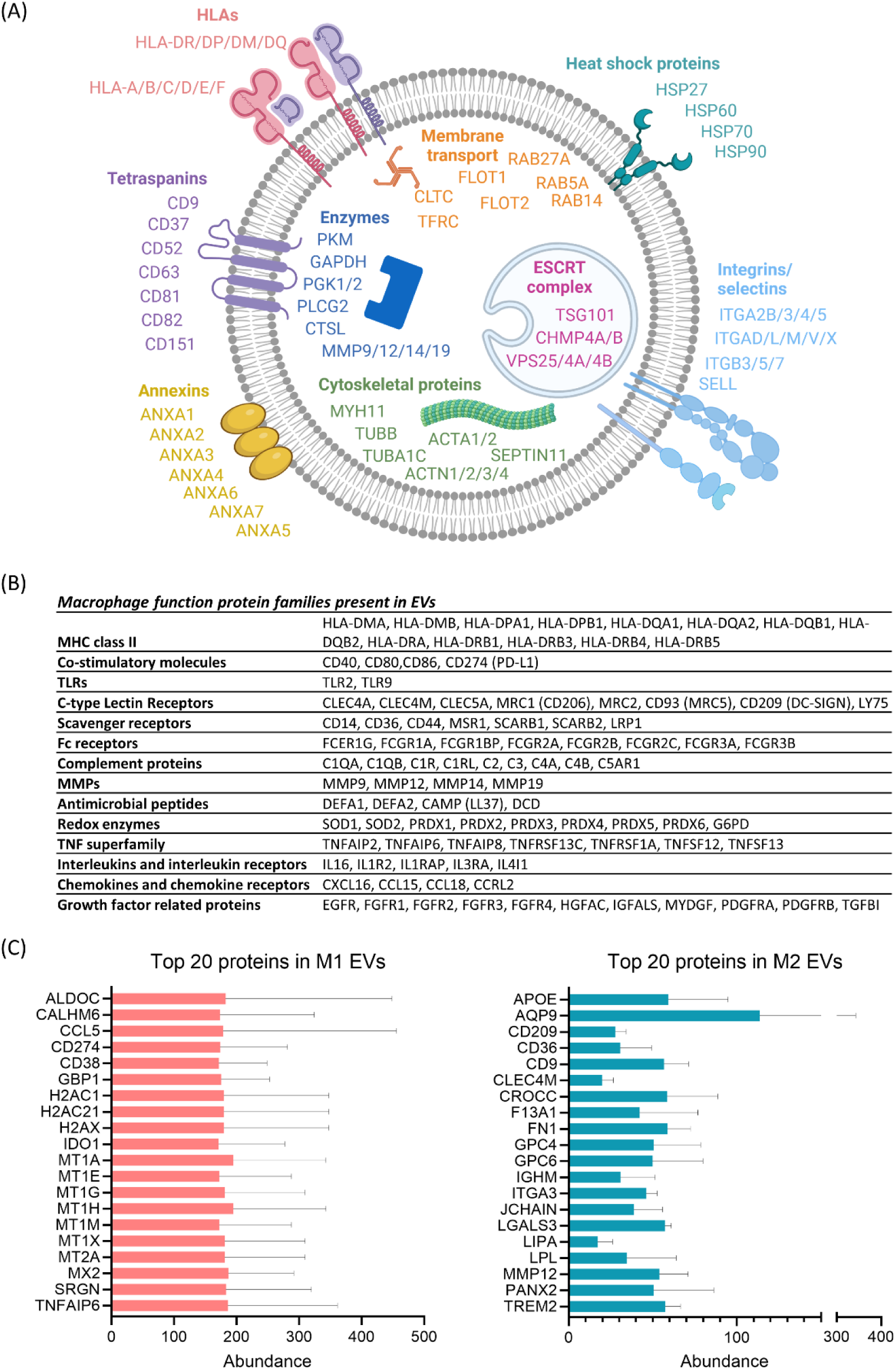
Proteomic characterisation of M1/M2 macrophage EVs. A) Proteins commonly associated with EVs present in M1 and M2 macrophage EVs. B) Proteins relevant to macrophage function present in M1 and M2 macrophage EVs. C) The top 20 most abundant proteins present in M1 (L) and M2 (R) EVs.

Prior to analysis to compare the protein cargo of M1 and M2 EVs, we first looked at the most abundant proteins detected in the EVs from these cells (Fig 2C). There was no overlap in the top 20 most abundant proteins in M1 and M2 EVs. Seven of the top 20 proteins in M1 EVs were metallothioneins (MT1A, MT1E, MT1G, MT1H, MT1M, MT1X, MT2A), a family of small, cysteine-rich proteins which bind metal ions. The others included proteins involved in macrophage activation, in line with their cellular source (CD38, CD274/PD-L1, CCL5/RANTES, TNFAIP6 and IDO1), intracellular signalling and defence (GBP1, MX2, CALHM6 and ALDOC), and chromatin structure and DNA damage response (histones H2AC1, H2AC21 and H2AX). CD38 is a cell surface glycoprotein involved in macrophage activation and proliferation. It’s expression is upregulated with exposure to lipopolysaccharide (LPS), used to polarise cells to M1 macrophages, and knockout or blockade of CD38 inhibits LPS-induced M1 polarization of macrophages [12]. PD-L1/CD274 is a transmembrane protein involved in supressing CD4/CD8 T cell activation and proliferation. It’s expression is induced by LPS treatment of monocyte/macrophages[13], as well as by IFN-γ, both used for the polarisation of our cells to M1 macrophages [14].

The top 20 most abundant proteins in M2 EVs included APOE (apolipoprotein E), several cell surface receptors and transporters (CD209/DC-SIGN, CD36, CD9, CLEC4M AND TREM2), enzyme and enzyme inhibitors (fibronectin 1, glypican 4/6 and galectin 3), proteins involved in ion and water transport (AQP9 and PANX2) proteins involved in adhesion and extracellular matrix interactions (FN1 (Fibronectin 1), GPC4 (Glypican 4), GPC6 (Glypican 6), LGALS3 (Galectin-3)), and immunoglobulin components (IGHM and JCHAIN). CD209/DC-SIGN is a C-type lectin found on the surface of macrophages that binds to high-mannose type N-glycans found on bacteria, viruses and fungi. CD209 is often reported as a marker of M2 macrophages, and we previously reported that our source cells have a 9-fold increase in CD209 mRNA compared to M1 macrophages [10]. Galectin is a pleiotropic lectin, including roles in cell growth and differentiation, cell adhesion, chemotaxis, apoptosis, and responses to bacteria and fungi. It has been implicated in both pro-inflammatory and pro-resolution responses. In RAW264.7 macrophages, it’s overexpression and secretion is associated with M2-type macrophages, whereas its downregulation promoted polarization to M1-type macrophages [15].

In a previous quantitative proteomics study of monocyte-derived macrophage EVs in response to β-glucan, the presence of immunity-related receptors, including high-affinity IgE receptor γ chain (FCER1G), C5a anaphylatoxin chemotactic receptor (C5AR1), cation-dependent mannose-6-phosphate receptor (M6PR), macrophage scavenger receptor (MSR1), and P2X7 receptor (P2RX7) in EVs was reported [8]. In our dataset of M1 and M2 monocyte-derived macrophages, we confirm the presence of FCERG1, M6APR, MSR1 and P2X7 in macrophage EVs, as well as a number of additional Fc receptor (FcR) fragments (FCER1G, FCGR1A, FCGR1BP, FCGR2A, FCGR2B, FCGR2C, FCGR3A, FCGR3B) and a range of additional scavenger receptors (CD14, CD36, CD44, MSR1, SCARB1, SCARB2, LRP1). Mast cells can release EVs bearing FcεR1 that bind to free IgE, reducing serum levels of IgE in an allergic mouse model [16], thus the presence of multiple Fc receptor proteins presents the possibility that macrophage EVs may also play a role in regulating serum or tissue immunoglobulins.

### Differentially abundant proteins in EVs from M1 and M2 macrophages

The Principal Component Analysis (PCA) plot (Figure 3A) illustrates the distribution of samples in the space defined by the first two principal components (PC1 and PC2), which together explained 47.86% of the total variance. M1 and M2 EVs form discrete clusters, indicated with ellipses, indicating distinct expression profiles. Comparison of M1 and M2 EV protein cargo identified 132 differentially abundant proteins (fold change >2, adjusted p value <0.05, peptide count >1; Figure 3B). Of these, 90 proteins were significantly increased in M1 EVs (Figure 3B; supplementary table 2), and 42 proteins were significantly increased in M2 EVs (Figure 3B; supplementary table 2). Figure 3C shows a heat map of the top 30 differentially abundant proteins in each EV type Looking at the top 10 differentially abundant proteins, M1 macrophages, typically involved in pro-inflammatory responses, exhibit highest upregulation of proteins associated with the stress response (SLC7A11, TXN, NAMPT), antiviral response (GBP1, GBP5, ISG15, STAT1, GBP5, MX2) and immune activation (GBP1,GBP5, TNFAIP2, CD38, ISG15, STAT1). Many of these have been previously associated with the M1 macrophage phenotype. Both GBP1 and GBP5 (Guanylate Binding Protein 1/5), together with ISG15 (ISG15 Ubiquitin Like Modifier), are associated with interferon (IFN) signalling. GBP1/5 are upregulated with high magnitude upon IFN-γ exposure [17], confer immunity against a range of intracellular bacteria [17], restrict a range of viruses [18], and their release within EVs has been described from monocyte-derived dendritic cells [19]. ISG15 is induced mainly by type I IFNs (IFNα/β), and has been previously reported in monocyte-derived macrophages [20]. Whether these proteins could exert anti-microbial functional effects in recipient cells, or could be a marker of viral infection, remains to be determined.

**Figure 3.**
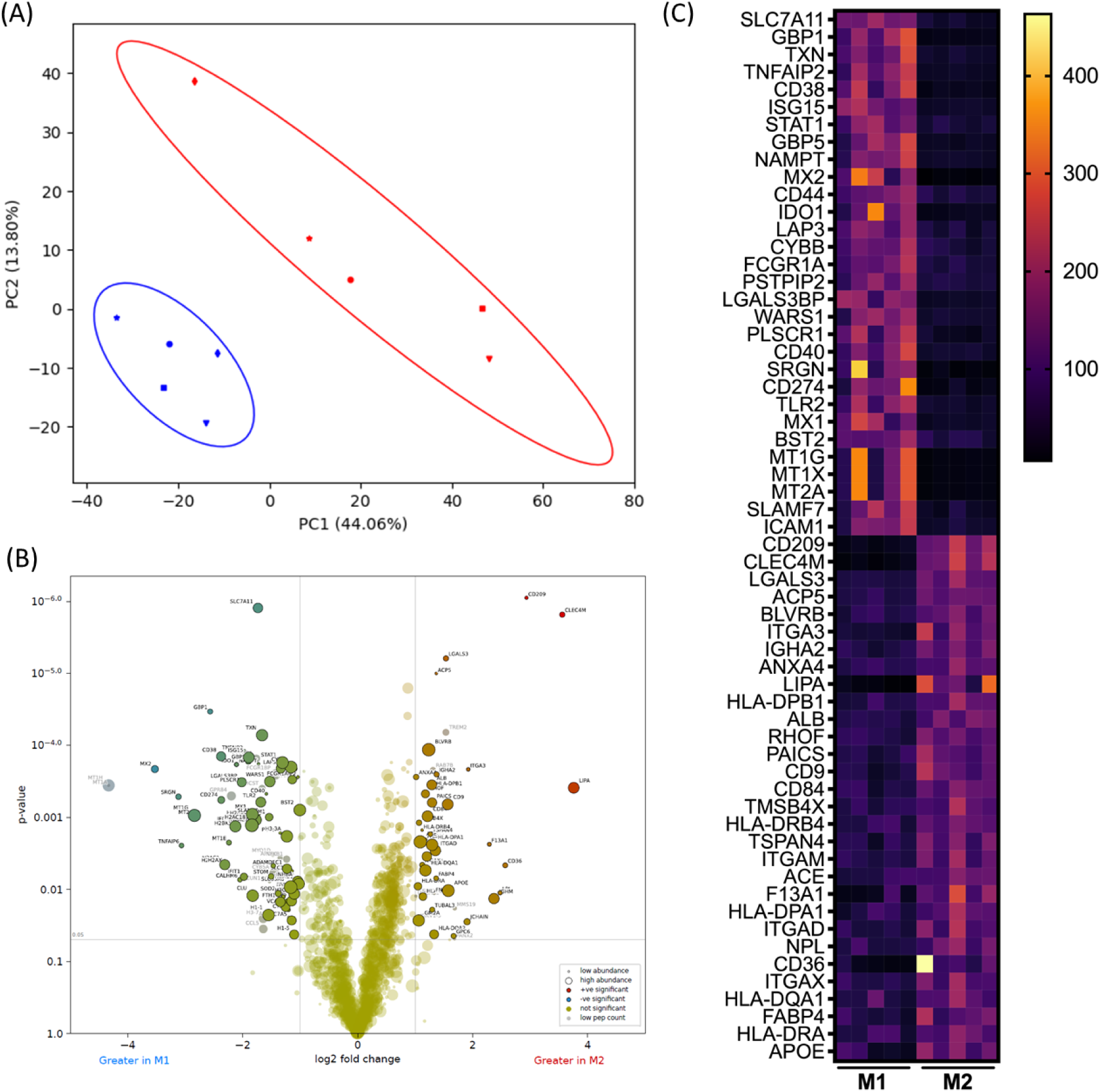
Comparison of protein cargo of EVs from M1 and M2 macrophages. (A) Principal component analysis (PCA) based on all proteins detected in M1 and M2 EVs. Plots show the first two principal components and their relative contribution to overall variance. Red data points are M1 EVs and blue are M2 EVs, and shapes represent the donor. (B) Volcano plot showing differential expression of proteins in M1 EVs (L) and M2 EVs (R). Coloured dots represent proteins that are significantly differentially abundant with a Log2 fold change greater/less than one and adjusted (FDR corrected) *p*-value < 0.05. The size of the dots reflects the peptide count. (C) Heatmap showing the top 30 differentially abundant proteins in M1 and M2 EVs (based on adjusted p value). Each row is a protein; columns represent the scaled abundance in the five biological replicates. Yellow-orange, higher abundance; purple-black, lower abundance.

Conversely, M2 macrophages, which are involved in anti-inflammatory responses and tissue repair, show highest upregulation of proteins associated with pathogen recognition, cell adhesion, and metabolic processes (e.g., CD209, CLEC4M, LGALS3, ACP5, BLVRB, ITGA3, IGHA2, ANXA4, LIPA, and HLA-DPB1). CD209/DC-SIGN is a C-type lectin that recognises high-mannose type N-glycans commonly found on bacteria, viruses and fungi, and is a commonly utilised marker of M2 macrophages [21]. Closely related to CD209/DC-SIGN in both structure and function, CLEC4M/L-SIGN is involved in cell adhesion and pathogen recognition [22,23]. LGALS3 is also a lectin but part of the galectin family (Galectin 3), and is involved in broader cellular functions including the immune response [24]. ACP5 (Tartrate-resistant acid phosphatase), BLVRB (Biliverdin reductase B) and LIPA (Lipase A, Lysosomal Acid Type) are all enzymes, involved in different metabolic processes (bone resorption, heme metabolism, lipid metabolism).

### Ingenuity pathway analysis demonstrates association of M1 and M2 EV cargo with canonical pathways relevant to infection and inflammation

To gain insight into the biological significance of the differentially abundant proteins, we performed Ingenuity Pathway Analysis (IPA) using the list of proteins that were significantly differentially abundant in M1 and M2 EVs. Of these, 77 proteins were eligible for IPA analysis for M1 EVs, and 39 for M2 EVs. There was a clear distinction in the biological roles of M1 and M2 EV protein cargo, aligning with the known function of their cellular source. The graphical summaries highlight that M1 EV protein cargo are associated with pathways including antiviral and antimicrobial responses, the immune response of antigen-presenting cells, and pathways including IRF7, IRF3, RIGI, STAT1, and several cytokines/chemokines (IL1b, TNF, IFNG, IFNB1) (Fig 4A). The graphical summary of M2 protein cargo shows association with immune function (CSF1, CSF2, IL4, IL5), the activation and phagocytic functions of macrophages, cell adhesion, and lipid metabolism (FGF2, FN1, PPARG, and INSIG1).

**Figure 4.**
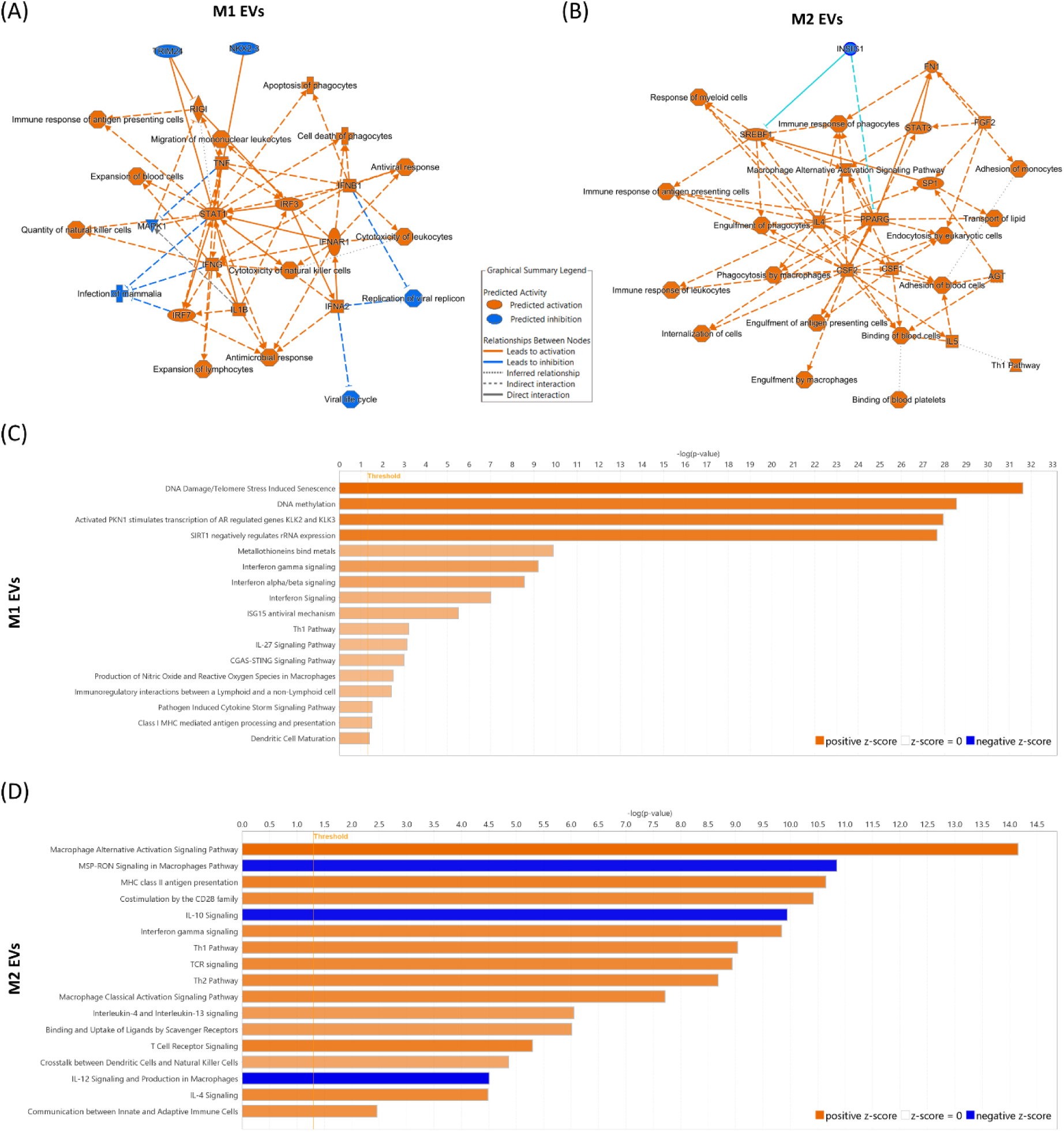
Graphical summaries and selected canonical pathways associated with M1 and M2 macrophage cargo. Values from the DESeq analysis were entered into a core analysis in IPA; one looking at proteins that were increased in M1 EVs, and one looking at proteins increased in M2 EVs (log fold change>1; adj p value<0.05). (A&B) Graphical summary overview of the major biological themes in the IPA Core Analysis of and how these relate to one another. Orange lines depict a relationship that leads to activation, and blue lines depict a relationship that leads to inhibition. (C&D) Selected canonical pathways of proteins upregulated in (C) M1 EVs and in (D) M2 EVs, focusing on immune-related pathways with z score>2 (full lists: Supplemental Table 3). Orange colour depicts predicted activation, and blue colour indicates predicted inhibition.

**Table 1 and Table 2** presents the top canonical pathways, diseases and disorders, and molecular and cellular functions derived from the data on M1 and M2 EVs, respectively. For M1 EV cargo, the top canonical pathways include DNA damage and methylation of DNA and histones, and regulation of rRNA. Diseases and disorders included several relevant to the immune response, including immunological disease and inflammatory diseases. Molecular and cellular functions included cellular death and maintenance, as well as cell-to cell signalling, in line with the EV compartment. For M2 EV cargo, the top canonical pathways include the macrophage alternative activation signalling pathway, in line with their source, as well as antigen/MHC class II antigen presentation and CD28 costimulation. Diseases and disorders are broad, including endocrine, gastrointestinal and metabolic diseases. Molecular and cellular functions again included cell-to-cell signalling, as well as cellular development, growth and proliferation, and lipid metabolism.

**Table 1.** Top canonical pathways and Diseases and Biofunctions derived from IPA core analysis of significantly increased protein cargo in M1 macrophage EVs.

| TOP CANONICAL PATHWAYS |  |  |
| --- | --- | --- |
| *Name* | *p value* | *overlap* |
| DNA Damage/Telomere Stress<br>Induced Senescence | 2.40E-32 | 32.1% 18/56 |
| DNA methylation | 2.80E-29 | 53.8% 14/26 |
| Activated PKN1 stimulates<br>transcription of AR regulated<br>genes KLK2 and KLK3 | 1.16E-28 | 50.0% 14/28 |
| SIRT1 negatively regulates rRNA<br>expression | 2.23E-28 | 48.3% 14/29 |
| PRC2 methylates histones and<br>DNA | 6.58E-27 | 40.0% 14/35 |
| TOP DISEASES AND BIO FUNCTIONS |  |  |
| Diseases and disorders |  |  |
| *Name* | *p value range* | *#Molecules* |
| Immunological Disease | 3.17E-03 - 1.47E-18 | 56 |
| Inflammatory Disease | 3.17E-03 - 1.47E-18 | 50 |
| Organismal Injury and<br>Abnormalities | 3.17E-03 - 1.47E-18 | 84 |
| Dermatological Diseases and<br>Conditions | 2.41E-03 - 9.17E-14 | 35 |
| Infectious Diseases | 3.17E-03 - 1.34E-13 | 46 |
| Molecular and cellular functions |  |  |
| *Name* | *p value range* | *#Molecules* |
| Cell Death and Survival | 3.17E-03 - 6.11E-12 | 40 |
| Cell-To-Cell Signaling and<br>Interaction | 3.17E-03 - 6.51E-10 | 29 |
| Cell Morphology | 3.15E-03 - 2.55E-09 | 22 |
| Cellular Development | 3.17E-03 - 3.26E-09 | 39 |
| Cellular Growth and Proliferation | 3.03E-03 - 3.26E-09 | 39 |

**Table 2.**
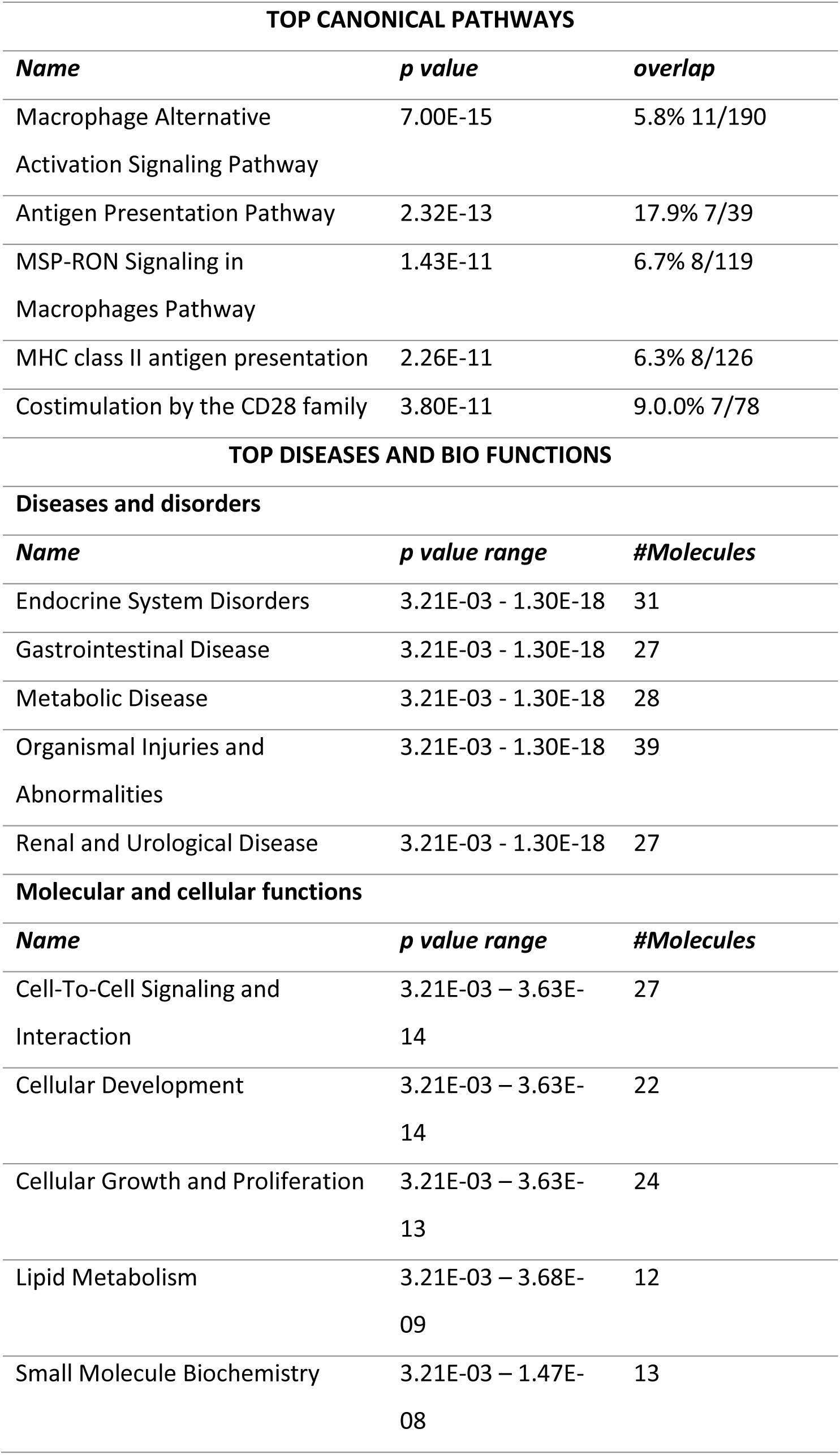
Top canonical pathways and Diseases and Biofunctions derived from IPA core analysis of significantly increased protein cargo in M2 macrophage EVs.

Further examination of canonical pathways related to M1 and M2 EV cargo revealed distinct associations. Full lists are shown in Supplemental Table 3, and selected pathways mainly linked to immunological responses are shown in Figure 4C-D. For M1 EV cargo (Figure 4C), pathways related to cytokine/chemokine signalling were abundant, particularly related to interferons. This pattern supports the role of M1 macrophages in initiating and sustaining pro-inflammatory responses and their potential involvement in immune responses against pathogens. The identification of pathways related to DNA damage, methylation, and oxidative stress suggests the involvement of M1 EVs in host defence mechanisms that manage pathogen invasion and the regulation of inflammatory responses.

In contrast, M2 EVs were associated with pathways related to immune modulation, including the macrophage alternative activation signalling pathway, antigen presentation, and lipid metabolism (Figure 4D). The activation of cytokine signalling pathways (IL-2, IL-4, IL-12, IL-13) and inactivation of the IL-10 signalling pathway align with the known roles of M2 macrophages in tissue repair, immune resolution, and regulation of metabolic homeostasis. The presence of lipid metabolism pathways (involving molecules like PPARG, INSIG1, and APOE) suggests that M2 EVs may play a role in lipid transport and metabolism, which could be critical for their role in maintaining homeostasis during the resolution of inflammation.

### IPA network analysis of M1 EV protein cargo

To gain insights into the molecular interactions and pathways associated with M1 and M2 EV protein cargo, we performed IPA network analysis on the differentially expressed proteins identified in our dataset. The analysis revealed 11 significant networks for M1 EV cargo (Supplemental table 4), with the top 5 networks summarized in Figure 5A. The top network identified involves molecules associated with Immunological Disease, Inflammatory Disease, and Organismal Injury and Abnormalities (Figure 5A-B). The network included molecules such as IFIT1, IFIT3, ISG15, STAT1, TRIM21, and various components of the interferon signalling pathway, highlighting its role in the body’s antiviral responses and the regulation of immune functions. ISG15 and STAT1 are key players in mediating these antiviral activities, while other molecules, such as JAK1/2 and MX1/2, underscore the network’s role in the broader interferon response. The presence of immunoglobulins (IgG2, IgG2b) and other immune-related proteins like SLAMF7 and TRIM21 further links this network to the immune system’s ability to respond to infections and inflammatory stimuli.

**Figure 5.**
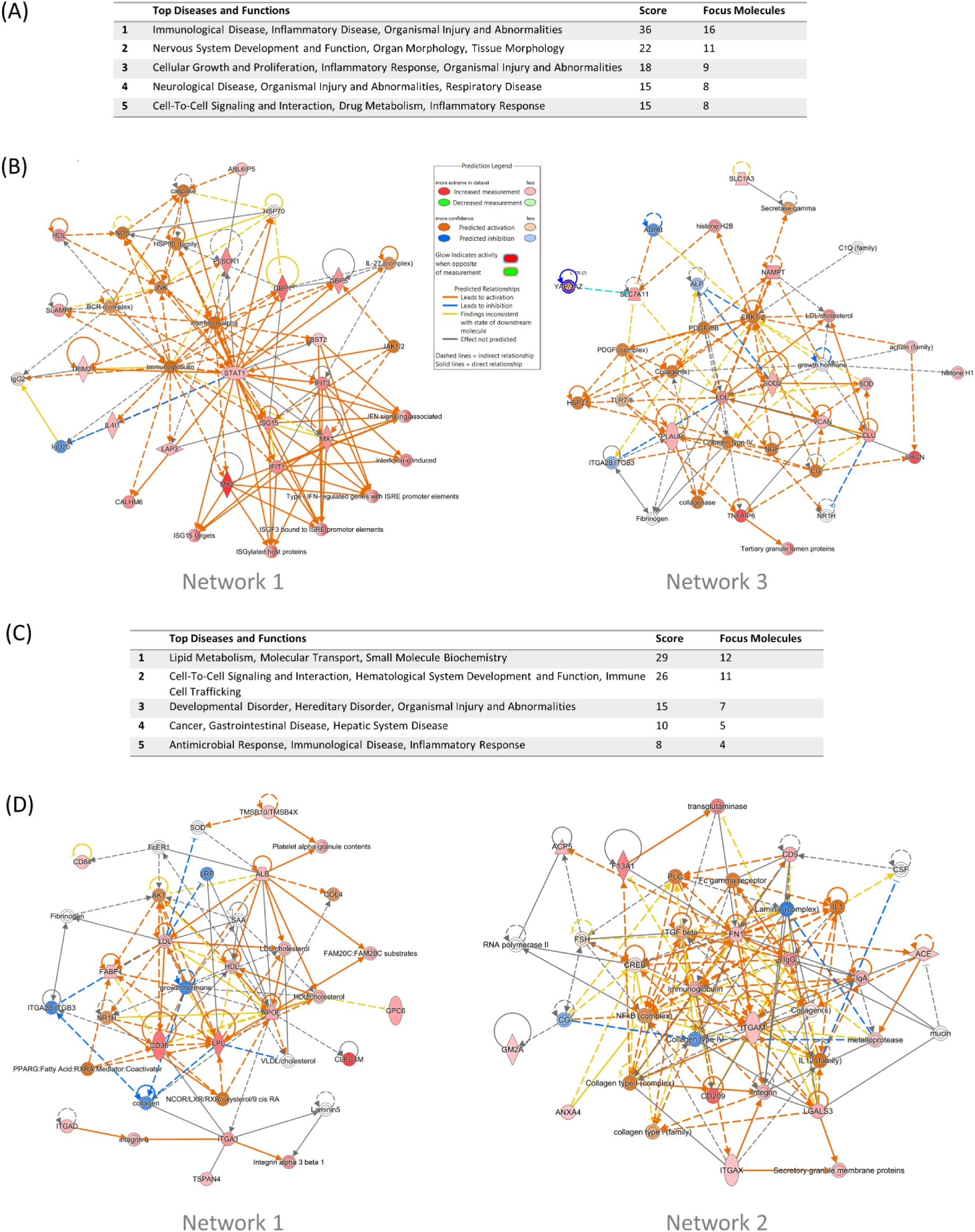
IPA network analysis of M1 and M2 EV protein cargo. (A) Table showing the top 5 networks in M1 EV protein cargo identified by Ingenuity Pathway Analysis (IPA). The five most significant functions for each network are listed, followed by the overall network score - a measure of the likelihood that the network represents a non-random set of molecules - and the number of molecules included in construction of the network (focus molecules). (B) Visualisation of networks 1 and 3 based on M1 EV cargo. Red nodes represent molecules present in the EVs, with intensity reflecting the level of expression. Orange nodes depict predicted activation, and blue depict predicted inactivation of other molecules in the network, respectively. (C) Table showing the top 5 networks in M2 EV protein cargo identified by Ingenuity Pathway Analysis. (IPA). The five most significant functions for each network are listed, followed by the overall network score, which is a measure of the likelihood that the network represents a non-random set of molecules, and the number of molecules included in construction of the network (focus molecules). (D) (B) Visualisation of networks 1 and 3 based on M2 EV cargo. Red nodes represent molecules present in the EVs, with intensity reflecting the level of expression. Orange nodes depict predicted activation, and blue depict predicted inactivation of other molecules in the network, respectively. The full lists, with associated molecules is shown in Supplemental Table 4.

The third most significant network involves processes related to Cellular Growth and Proliferation, Inflammatory Response, and Organismal Injury and Abnormalities (Figure 5A-B), suggesting that this network plays a role in response to damage and subsequent repair mechanisms. Network molecules included ERK1/2, PDGF-BB, LDL/cholesterol, TLR7/8, and members of the activin family. The presence of ERK1/2 and PDGF-BB highlights this network’s involvement in signalling pathways regulating cell growth, survival, and proliferation. Additionally, structural components like collagens and histones (H1 and H2B) suggest roles in maintaining cellular architecture and regulating gene expression.

### IPA network analysis of M2 EV protein cargo

IPA network analysis revealed five significant networks associated with M2 EV cargo (Figure 5C, Supplemental Table 4). The top network involved molecules central to lipid metabolism, molecular transport, and small molecule biochemistry (Figure 5C-D). Network molecules included AKT, APOE, PPARG, integrins, and various lipoproteins (LDL, HDL, VLDL), highlighting their interconnected roles in regulating lipid homeostasis, cellular interactions, and transport mechanisms. AKT and PPARG are important in mediating metabolic processes, while APOE and lipoproteins are essential for cholesterol transport. Integrins, which facilitate cell-extracellular matrix interactions, further underscore the network’s involvement in cellular adhesion and signalling.

The second network focuses on molecules integral to cell-to-cell signalling and interaction, haematological system development, and immune cell trafficking (Figure 5C-D). This network includes ACE, NFκB, integrins (e.g., ITGAM, ITGAX), collagen types I and IV, and key immune components like IL1, IL12, and immunoglobulins (IgA, IgG). These molecules interact to regulate immune responses, extracellular matrix integrity, and cellular communication. NFκB and integrins are central to immune signalling and cell adhesion, while collagens and laminin are vital for maintaining the structural framework of tissues. The presence of molecules like TGF-beta and metalloproteases highlights the network’s role in tissue remodelling and inflammatory responses.

### Metal homeostasis proteins are present and differentially abundant in M1 and M2 macrophage EVs

As seven of the top 20 most abundant proteins in M1 EVs represented metallothioneins - a family of proteins involved in metal homeostasis and oxidative stress responses – we assessed the presence of all metallothioneins in our dataset, and if other related proteins were present, and differentially abundant in M1 and M2 EVs. Nine metallothioneins were present in the dataset, and seven of these were significantly elevated in M1 versus M2 EVs (Figure 6A). Of note, networks 4 and 5 from the previous IPA network analysis of M1 EV cargo (Supplementary Table 4) involved several metallothioneins. Interestingly, metallothioneins are increasingly being implicated to play roles in infection and immunity [25]. In addition to metallothioneins, several other metal-binding proteins were identified in macrophage EVs; the light and heavy chain of ferritin (FTL and FTH1 respectively), transferrin (TF), and various haemoglobin subunits (HBA1,HBB, HBD, HBE1, HBG1, HBG2) (Figure 6B). Of these, the ferritin heavy chain FTH1 was higher in M1 EVs compared to M2. Several members of the solute carrier gene (SLCs) superfamily were present in macrophage EVs, including the copper transport SLC31A1 and the zinc transporter SLC39A8/ZIP8. The zinc transporter SLC39A8/ZIP8 was more abundant in M1 EVs. The metalloenzymes carbonic anhydrase (CA1, CA2, CA3), ceruloplasmin (CP), superoxide dismutases (SOD1, SOD2), catalase (CAT), and glutathione peroxidase (GPX1, GPX4) were all present, with SOD2 significantly higher in M1 EVs (Figure 6C). Alongside metallothioneins, many of these proteins are involved in oxidative stress response pathways. Additional proteins present also involved in oxidative stress responses included all six peroxiredoxins (PRDX1-6), and Glutathione S-transferase pi 1 (GSTP1).

**Figure 6.**
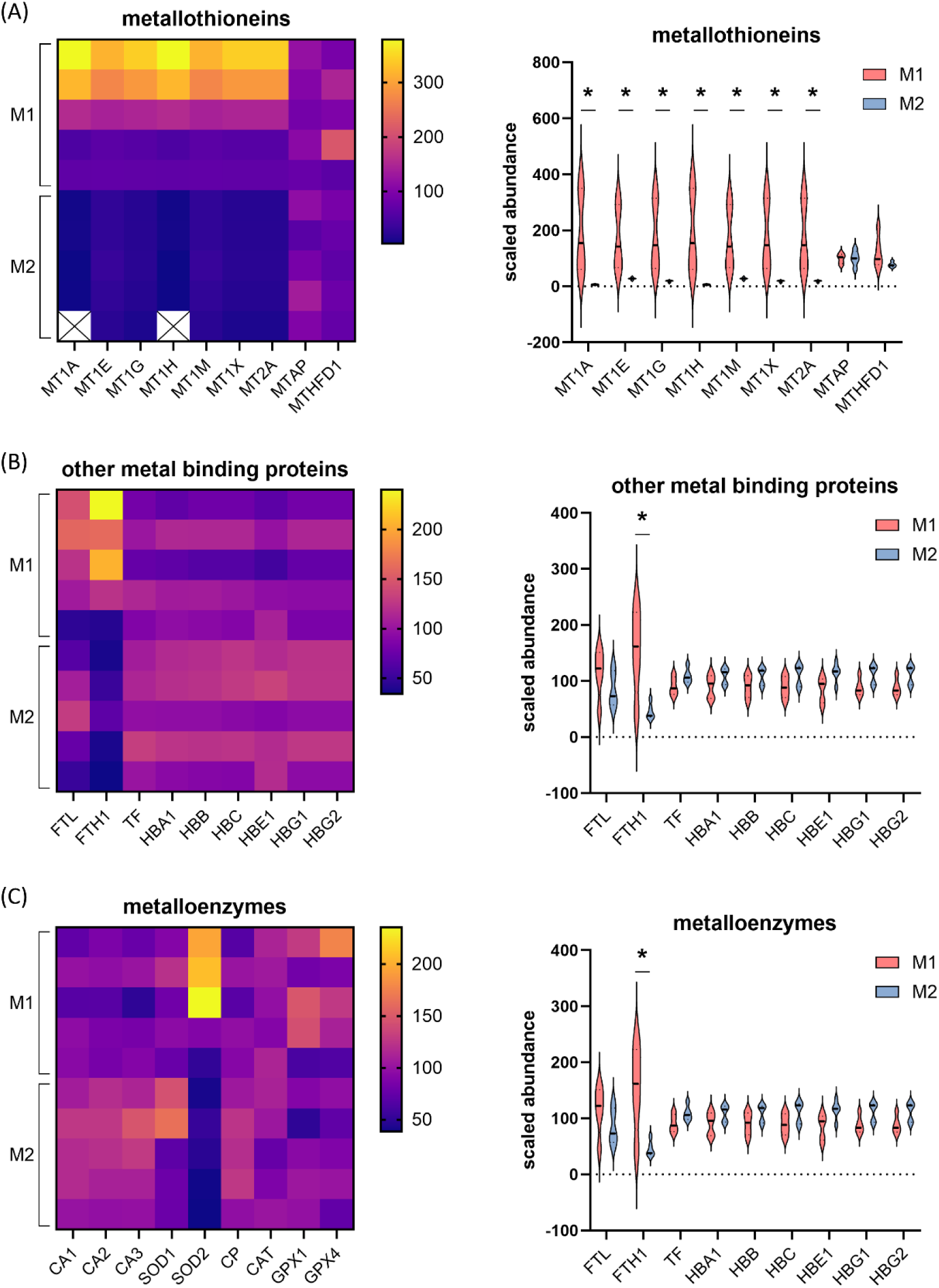
M1 and M2 macrophage EVs contain proteins involved in metal homeostasis and oxidative stress. Presence of (A) metallothioneins, (B) other metal-binding proteins, and (C) metalloenzymes in M1 and M2 EVs represented in heat maps and violin plots. * indicates significant differences between M1 and M2 EVs measured by DESeq (fold change>2 and adjusted p value<0.05).

## CONCLUDING REMARKS

Proteomics has previously been applied to the study of monocyte-derived macrophage EVs in response to pathogens and pathogen associated molecular patterns [7,8]. This current study reports the isolation and characterization of EVs from M1 and M2 monocyte-derived macrophages, the comprehensive protein cargo of these EVs, and differences in protein abundance between M1 and M2 EVs, representing a highly useful dataset for future investigations.

The strengths of this study includes the use of five biological replicates, well-characterised cells [10], entirely serum-free culture of source cells, and employment of TMT quantitative proteomics, which together resulted in our ability to identify a large number of proteins (>1,500; >2,000 including isoforms) in macrophage EVs. The weakness of this study is that we only generated EVs from macrophages from female donors, and so we are unable to explore sex differences in macrophage EV cargo, which would be an important area of future investigation. However, our reporting of biological sex is also a strength, as many proteomics studies do not include this information. Finally, as we did not perform proteomics on the cells producing the EVs, we were not able to analyse which EV proteins are a direct mirror of the source cell, and which are enrichment/depleted compared to the cells, which could indicate where there is preferential packaging of specific proteins into EVs.

The distinct proteomic profiles of M1 and M2 EVs presented here highlight their distinct cellular source, as well as suggesting that they may have specialized roles in modulating immune responses, with M1 EVs contributing to pro-inflammatory and antiviral activities, and M2 EVs supporting immune regulation and tissue repair. These findings have significant implications for understanding the complex interplay between different macrophage phenotypes and their roles in health and disease. The differential protein cargo of M1 and M2 EVs could serve as potential biomarkers for disease states characterized by dysregulated macrophage activity, such as chronic inflammatory diseases, infections, and cancer. Future studies should investigate the functional impact of these differentially abundant proteins in relevant disease models to further understand their roles in macrophage-mediated immunity. Finally, exploring the potential therapeutic applications of modulating EV cargo or their release from macrophages may provide novel strategies for treating immune-mediated disorders.

## ACKNOWLEDGEMENTS

This work was supported by NICHD, NIH (5R01HD093801, awarded to BH). This work was also supported by the National Institute for Health Research Comprehensive Biomedical Research Centre at Imperial College Healthcare NHS Trust and Imperial College London. The views expressed are those of the author(s) and not necessarily those of Imperial College, the NHS, the NIHR or the Department of Health. The authors wish to thank Simrit Sahota, Vladimir Bokun and Alice Hawkins for contribution to discussions. We also wish to thank Dr Olivia Raglan for assistance with IPA, Andrew McArdle for advice on proteomics analysis, and Prof. Wolfgang Maret for providing expertise regarding metallothioneins. The graphics were created with Biorender.

